# A practical method to improve the efficiency of pollination in maize breeding and genetics research

**DOI:** 10.1101/2023.04.04.535612

**Authors:** Dylan L. Schoemaker, Frank McFarland, Brian Martinell, Kathryn J. Michel, Lucas Mathews, Dan O’Brien, Natalia de Leon, Heidi F. Kaeppler, Shawn M. Kaeppler

## Abstract

Seed increase through manual pollination is a critical part of maize breeding and genetics research to advance generations in breeding programs, to create desired research crosses, and produce hybrid seed for trials. Pollination in the field and in controlled environments relies on the availability of high-quality pollen at the time that recipient silks are receptive. Generally, pollinations are made by capturing pollen from the tassel in a paper pollinating bag placed on the tassels one day prior to pollination and newly released pollen is then transferred to silks on the target plant. In the field, maize pollen is only viable for one to four hours following dehiscence and the rate of desiccation is influenced by environmental conditions. We have developed a method which increases the lifespan of pollen and allows pollen from a single tassel to be used to pollinate many ears by mixing fresh pollen with a dilutant that can be stored for multiple days. We identified characteristics of the size of suitable substrates and selected a PEEK based substrate for regular utilization. We evaluated pollen viability and empirically demonstrated the capability to store pollen up to nine days when pollen is mixed with a PEEK substrate and stored at 6°C. The pollen storage method was used to make successful pollinations across 24 maize inbred lines tested and was generally equivalent to the standard manual pollination process. This method has the potential to increase the efficiency of breeding operations and may be useful in an array of genetic studies.

**Core Ideas:**

• Manual pollinations in breeding and genetics research requires pollen available when recipient silks are viable.
• The method collects and stores maize pollen for at least five days and facilitates efficient pollination.
• Pollen is mixed with polyetheretherketone and uses field-collected pollen and simple storage conditions.
• The method can increase the number of pollinations per tassel and generates a reasonable number of viable seeds.

## INTRODUCTION

Access to a sufficient quantity of high-quality pollen when silks of target plants are receptive is vital for seed production associated with maize breeding and genetics research. Maize pollen is generally short-lived and sensitive to extreme moisture and temperature (Jones & Newell, 1948; Barnabas, 1985; Buitink et al., 1996; Luna et al., 2001). Methods to store pollen for later use, and to increase the efficiency of the pollination process, would provide a substantial benefit to plant breeding and genetics research.

Pollen storage and viability has been studied by researchers since the early 1920s to aid breeding and genetics research (Anthony & Harlan, 1920; Knowlton, 1922). Some of these early studies have shown that pollen longevity varies across species. For example, barley (*Hordeum vulgare*) pollen exposed to free air for 10 minutes was inviable due to moisture loss (Anthony and Harlan, 1920). Alternatively, potato (*Solanum tuberosum*) crops are considered desiccation tolerant (Towill, 1981) because pollen remains viable after desiccation to moisture content as low as 5% to 7% (Roberts, 1973). Further, Kesseler (1930) reported that potato pollen can be viable after 14 days with minimal storage treatments if kept at 15% to 20% relative humidity. When potato pollen was stored at -20°C for 11 months, the stored pollen generated as many seeds as fresh pollen (Howard, 1958). Pine (*Pinus ponderosa*) is also desiccation-insensitive and displays a faster rate of pollen-moisture loss relative to maize when placed on MgCL_2_ or Mg(NO_3_)_2_ (Connor & Towill, 1993).

Differences among species in the rate of pollen-water loss can affect long-term pollen storability. For example, broccoli (*Brassica oleracea* var. *italica*) pollen stored in liquid nitrogen for two months resulted in 43% germination success (Crisp and Grout, 1984). Alternatively, *Linum longiflorum* and maize pollen stored for five months at 0°C to 5°C led to a 25% and 15% pollen germination rate, respectively (Nath & Anderson, 1975).

Beyond storing pollen for seed generation, collecting pollen prior to dehiscence can help minimize unintended gene flow and therefore contribute to the development useful genetics materials for research. In maize for example, genetically modified (GM) pollen can be blown by the wind into neighboring fields and lead to genetic erosion (Rogers & Parkes, 1995; Serratos et al., 1997). As maize pollen is blown via the wind, isolation nurseries are needed to minimize gene flow from aerial pollen. However, the effective isolation distance is a function of windspeed, direction, and circulation (Bateman, 1947a and 1947b; Jones & Brooks, 1950; Raynor et al., 1972; Luna et al., 2001).

Minimizing off target pollen movement is also critical for maize hybrid seed production to ensure purity of hybrid cultivars. Maize hybrid seed production relies on the large quantities of windblown pollen from one inbred line landing and germinating the receptive stigma of an adjacent inbred (Heslop-Harrison, 1979; Kiesselbach, 1999). However, this system is resource intensive and seed production yield decreases when the anthesis-silking interval expands beyond three days and/or an inbred line has a narrow pollen shed window (DuPlessis & Dijkuis, 1967; Wych, 1988; Arisnabarreta & Solari, 2017). However, the risk of these latter issues can be minimized via efficient methods for collecting and dispensing stored pollen. PowerPollen has developed the first proprietary system for bulk collection, preservation, and on-demand application of stored maize pollen via electronic sensors attached to a distribution apparatus on a tractor (Cope & Krone, 2016). The technology and protocols increase seed production yield up to 40% and allows breeders to select inbred parents with greater flexibility (https://powerpollen.com/corn-seed/).

When maize pollen is collected, it must be quickly transferred to a substrate to avoid desiccation as maize pollen is short lived (Berjak et al., 1992). Common substrates previously used for storing pollen include organic solvents (Iwanami & Nakamura, 1972), polyethylene products, and chemical treatments. Barnabas and Rajki (1976) described the use of a polyethylene substrate for maize pollen storage. Mineral oil is another substrate used to manipulate pollen. For mutagenesis, mineral oil is mixed with EMS and applied to fresh maize pollen as a chemical treatment. The treated pollen is then used to pollinate plants with receptive silks to produce mutagenised offspring (Neuffer & Coe, 1977; Settles, 2020).

Beyond identifying an appropriate substrate, the relative moisture content of the pollen and ambient temperature were initially shown to influence storability of maize pollen. Once the pollen and substrate are mixed and placed in an airtight vessel, the container can be kept in liquid air (Collins et al., 1973) or nitrogen at -192°C or -196°C, respectively, for long-term storage. Barnabas et al. (1988) further demonstrated that when maize pollen is stored at low temperatures in liquid nitrogen, a 13% pollen water content was optimal for storing pollen up to one week after pollen collection and led to a 78% seed set.

Deep-freezing storage methods can potentially maintain pollen viability for up to a year. Maize pollen mixed with a polyethylene-based substrate placed in a sealed vessel generated viable pollen granules after a year of storage (Barnabas & Rajki, 1976), while soybean pollen-maintained viability for four months if kept at -20°C (Tyagi & Hymowitz, 2003). While these deep-freezing techniques are effective at supporting pollen viability for long term storage, Jones and Newell (1948) focused on cost effective techniques for short term storage. Seed set from stored maize pollen was observed after 48 hours of storage and pollen viability was maintained up to eight days if kept at 4.4°C and 90% relative humidity (RH) but decreased to six days if RH decreased by 10% (Jones & Newell, 1948). These results suggest that maintaining proper RH is important for minimizing maize pollen grain desiccation during short term storage.

Maize pollen is short lived due to rapid pollen-water loss following dehiscence (Jones & Newell, 1948; Barnabas 1985; Buitink et al., 1996; Luna et al., 2001). External factors such as humidity, wind, and temperature can accelerate water loss (Roy et al., 1995; Schoper et al., 1987a; Schoper et al., 1987b) and limit viable pollen availability during seed production. Compared to other species, maize is considered desiccation intolerant as viability dramatically decreases when pollen water-content is below 0.4 g g^-1^ (Buitink *et al.,* 1996). Luna *et al*. (2001) used *in vitro* pollen germination assays to demonstrate that pollen could survive for two hours following dehiscence when released from maize plants grown in an environment with average daily high temperatures ranging from 28°C to 30°C and average RH from 31% to 53%. However, pollen viability was influenced by atmospheric water potential (Luna et al., 2001). Pollen drift will vary by location as the pollen grain temperature will match the air temperature of a given environment (Aylor, 2003). These results were further supported by Aylor (2004), who observed a 50% reduction in maize pollen germination after pollen was exposed to direct sunlight and air for 60 to 240 minutes.

The goals of this study were to develop and empirically evaluate methods that would permit cost-effective short-term maize pollen storage under practical field conditions and facilitate increased efficiency of pollination in breeding and genetics research. We evaluated different storage method across multiple field-based settings and different genetic backgrounds. We have utilized this technique extensively in our research program and have found it to be reliable and to increase pollination process efficiency.

## 2 MATERIALS AND METHODS

### 2.1 Storage substrate identification

Five potential storage substrates were initially tested to mix with maize pollen. These included Aeroperl 300/30 from Evonik (product code: 10024572), Sipernat 22 S from Evonik (product code: 99002421), Sipernat D 13 from Evonik (product code: 10020326), Blue Polyethylene Microspheres (PEM) from Cospheric (product line: BLPMS-1.00), and DicaLite^(R)^ Natural Diatomaceous Earth from Dicalite Management Group. Each medium was mixed with pollen collected from maize inbred line PHAJ0 grown in a seed production nursery at the West Madison Agricultural Research Station in Verona, WI during the summer of 2019. Pollen was collected by removing tassels pre-pollen shed and placing them in a FloraLife Crystal Clear^(R)^ Flower Food 300 liquid medium under cool-white T12 fluorescent lights to promote anther exertion. When 50% of the tassel was shedding pollen, anthers were shaken off the tassel branches and placed into a 120 ml (4 oz) sterile cup. First, the anthers and large debris were removed by sieving the pollen through a stainless-steel strainer to remove anthers and large debris (Figure 1B). The pollen was then sieved again through a size 80 mesh (0.180 mm) using a Tansoole Experimental Sieve to remove small clumps of pollen (Figure 1C). The sieved pollen was then independently mixed with each of the five storage substrates at a ratio of one part pollen to five parts substrate (1:5) and poured into a glass scintillation vial (Figure 1D and Figure 1F). The substrate and pollen mix were held horizontally and gently rotated approximately five times until the medium and pollen was homogenized (Figure 1E). The mixture was either kept in a sealed 120 ml (4 oz) sterile sample cup and stored in a walk-in cold-room at 4°C (Figure 1F) or directly used to pollinate plants with receptive silks (Figure 1G).

**Figure 1.**
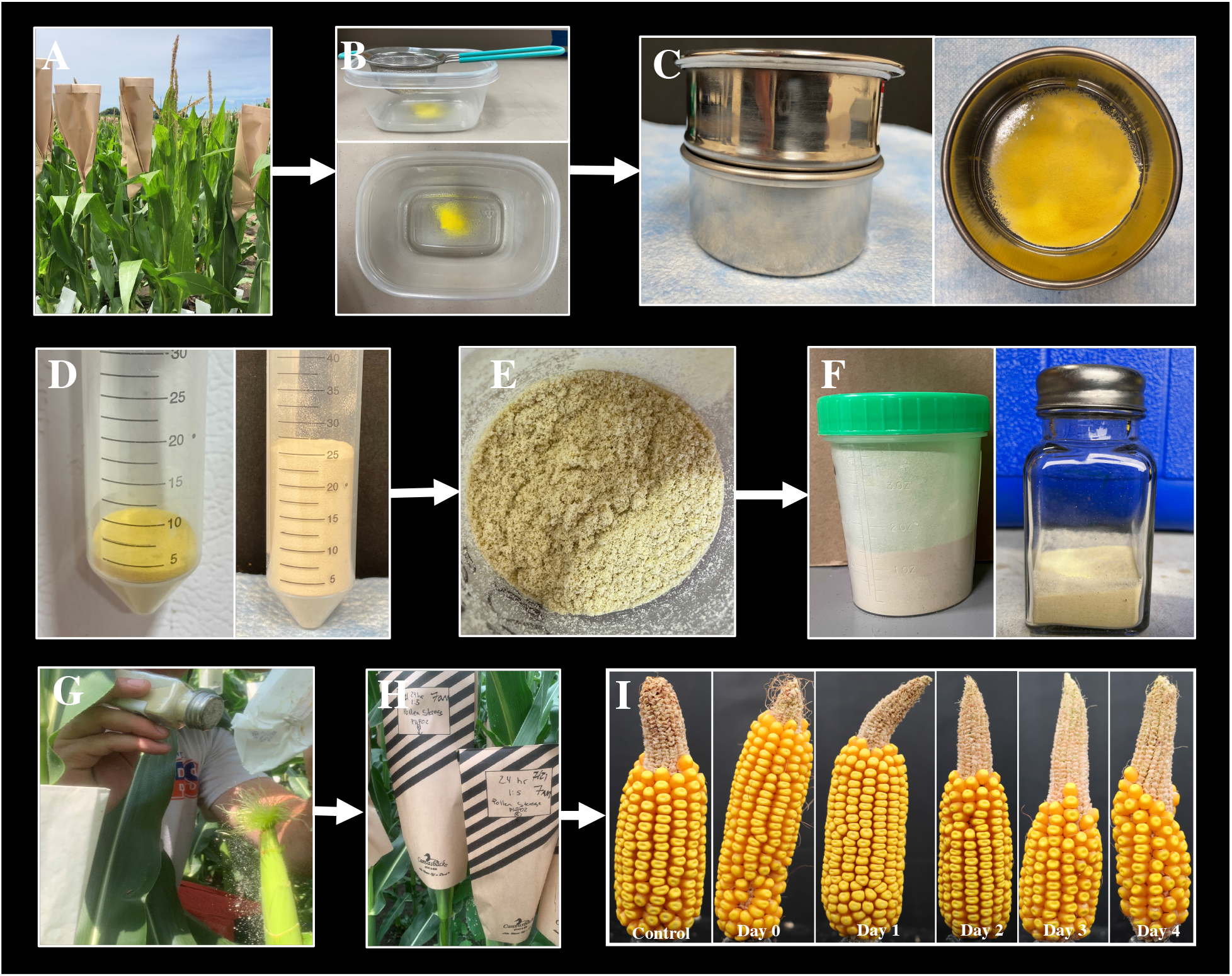
Flowchart describing the process of pollen collection, storage, and application. A) Tassel bags placed on the inflorescence of the pollen parent 24 hours in advance of pollen collection are removed and the pollen B) is dumped through a metal strainer to remove large debris and a C) 0 mesh sieve is used to remove clumped pollen. D) A concentration of one part pollen to five parts (1:5) PEEK-MP140 is used and E) mixed. F) The mixed pollen is immediately stored between 4°C and 6°C in a sealed tight container or transferred to a glass spice container for application on plants with receptive silks. G) Approximately 0.047 g (±0.003) of mixed pollen is applied per ear shoot and H) pollinated ears are covered with a tassel bag. I) Examples of ears pollinated with mixed pollen compared to a control self-pollination (far left) when mixed pollen is stored out to four days.

The mixed maize pollen was applied to ear shoots of seed parent inbred lines that were covered prior to silk emergence to ensure ovules were pollinated from stored pollen and to prevent contamination from adjacent plants. To make pollinations, the pollen mixture was gently rotated three times, and a small portion of the dilution was aliquoted into an application vessel that was either a 50 mL falcon tube or a 2.7 oz glass spice container with approximately five to ten one-millimeter diameter holes. Approximately three ‘shakes’ of mixed pollen from the vessel was applied to each ear where a ‘shake’ is defined as the movement of the applicators arm from a 90° to 45° angle when the container is maintained perpendicular to the forearm (Figure 1G). Based on the average of 20 replicates, approximately 0.047 g (±0.003 standard error) mixture of pollen-substrate is applied per maize ear. A tassel bag was immediately placed over the ear shoot following pollination and stapled together on the opposite side of the ear to prevent pollen from adjacent plants landing on the inbred silks (Figure 1H).

Each of the pollen mixtures were applied to two plants of maize inbred line LH244 with receptive silks after 2, 6, and 21 days of storage between approximately 9:00 A.M. and 11:00 A.M. On the same day, undiluted stored pollen from PHAJ0 was applied onto two LH244 plants with receptive silks as a control. Ears were directly covered after the application and harvested two weeks later. The number of kernels on the each of the four ears was visually counted at the time of harvest.

An additional storage medium, PEEK-MP140, manufactured by PolyClean Technologies Inc., was evaluated using a field setting at the West Madison Agricultural Research Station in Verona, WI during the summer of 2020. PEEK-MP140 is a fine milled powder made from recycled Polyetheretherketone (PEEK), 450G. Pollen was collected, stored, and applied to targeted plants with receptive silks using the method described above and in Figure 1. Supplemental File S1 lists all the steps used to collect, make, and apply mixed pollen to plants with receptive silks. For evaluation of PEEK-MP140, pollen was collected by placing a tassel bag on the inflorescence of the pollen parent 24 hours in advance and collecting freshly released pollen in the bag. The collected pollen in the tassel bag was sieved through a metal strainer and size 100 mesh (0.154 mm) to remove anthers and large debris prior to mixing (Figure 1B). The utility of PEEK-MP140 as a storage substrate was evaluated by storing both a one-part pollen to five-part substrate (1:5) and a one-part pollen to ten-part substrate (1:10) mixture to evaluate how the concentration of pollen influences grain fill. The mixture was stored in a walk-in cold room at 6°C and then used to pollinate five different plants of a commercial inbred line with receptive silks every day at mid-morning for eight days.

### 2.2 Scanning Electron Microscopy imaging

Both PEEK-MP140 and Cospheric blue polyethylene microspheres were further analyzed using scanning electron microscope (ESM) at the Wisconsin Newcomb Imaging Center (NIC). All high-resolution images of maize pollen within the medium were captured on a FEI Quanta 200 microscope set to low vacuum (ESEM mode). Prior to imaging, pollen was collected from inbred line LH244 grown in a greenhouse at the Wisconsin Crop Innovation Center (WCIC) in Middleton, WI by placing a tassel bag on the inflorescence 24 hours prior to pollen collection. After 24 hours, the fresh pollen was collected, sieved, and mixed with PEEK-MP140 and PEM at a 1:5 ratio, as previously described. The mixture was stored for 24 hours at 6°C in a standard refrigerator prior to imaging.

### 2.3 Experimental design of field trials

The utility of stored maize pollen for breeding and genetics research was assessed using field settings during the summer of 2020, 2021, and 2022 at the West Madison Agricultural Research Station in Verona, WI. Pollen was collected from inbred lines grown in 12 ft long, single-row plots using the methods described above. The pollen was sieved and diluted with medium at a station adjacent to the field using the procedure described in Figure 1 and Supplemental File S1. The pollen-substrate mixture was either directly transferred to a 2.7 oz glass spice container (Figure 1F) and applied to plants with receptive silks (Figure 1G) in the mid-morning or kept in a sealed airtight container and stored at 6°C in a walk-in cold-room for later application.

### 2.4 Experimental assessment of stored pollen over time

In 2020, the method and substrate for storing maize pollen was initially tested by collecting pollen from a line heterozygous for purple pigmented kernels and applying it to ears of plants that did not have pigmented aleurone or endosperm. Pollen from the purple kernel inbred line was collected and stored at 6°C in a walk-in cold-room from one to eight days and mixed with PEEK-MP140 at both a concentration of 1:5 and 1:10. For each of the eight days, five pollinations were made between approximately 8:00 A.M and 10:00 A.M. After approximately 40 days after pollinations (DAP), ears from all five replicate pollinations per pollen concentration and days of storage treatment were collected from the seed parent and visually inspected to determine if kernels were present on the ear. The proportion of ears out of the five replicate pollinations per treatment with at least 10 kernels was recorded.

In 2021, an experiment was conducted to evaluate how the ratio of pollen to substrate affected grain fill and determine if the time-of-day mixed pollen is applied to receptive maize silks impacts seed set. Pollen was collected from the maize inbred PHP02 and mixed with PEEK-MP140 as previously described. The mixed pollen was then stored for up to 48 hours in both a 1:5 and 1:10 dilution. Each day, both mixtures were used to pollinate six plants with receptive silks of PHP02 every hour between 7:00 A.M. and 12:00 P.M. The ears pollinated with stored pollen were harvested between 35 and 45 DAP and two images of each ear were captured as previously described. Grain fill was assessed using the images by visually rating the two images per ear for the proportion of the ear filled with grain on a one to ten scale (Supplemental Figure S1) and assigning each ear an average grain fill rating based on the two images.

The average percent grain fill over the six replicate pollinations was analyzed using an analysis of variance (ANOVA) to test for the effect of the timing of the pollen application and pollen to substrate ratio using the equation y_ij_ = Time_i_ + Ratio_j_ + ε_ij_. Time refers to the effect of the i^th^ time between 7:00AM and 12:00P.M. and Ratio corresponds to the effect of j^th^ pollen to substrate ratio being either 1:5 or 1:10. The residuals were independent and identically distributed, 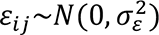. A Tukey Honest Significant Difference (HSD) test was conducted *post-hoc* using an experimental wise error rate (#x03B1;*_E_*) of 5% to test for significant differences between each combination of time and ratio.

In 2022, grain fill from stored pollen was studied across two different fields planted on May 11^th^ and June 3^rd^, corresponding to an early and late planting date for our region, respectively. Pollen from the maize inbreds LH244, LH287, and PH24E was collected from tassels when at least 50% of the plant’s main tassel was shedding pollen. The pollen was mixed with PEEK-MP140 at a ratio of 1:5 and stored up to 10 days at 6°C. The pollen mixture for each inbred line was used to pollinate six plants with receptive silks of LH244 each day, including the initial day of collection (Day 0). Each day, an additional three self-pollinations were made using the standard bagging method as a control by taking pollen directly from a tassel bag that was placed the previous day on the inflorescence of the seed parent inbred LH244 and directly transferring the pollen to the ear.

The ears pollinated with the stored mixed pollen were collected between 40 and 45 days DAP. For each ear, an image was captured. The ear was then rotated 180° and second image was recorded such that there were two images per ear. A visual rating for percent grain fill was given to each image based on a one to ten scale (Supplemental Figure S1) and the number of kernels on the ear were visually counted. The average visual rating across the two images and total kernel count across the two images per ear was used for further analysis.

The average number of kernels per ear and average percent grain fill over the six replicate pollinations was analyzed using an ANOVA in R-software based on the model y_ij_ = Storage_i_ + Planting_j_ + ε_ij_. Storage refers to the i^th^ number of days that the pollen mixture was stored prior to making pollinations in the field. The terms Inbred and Planting refer to the effect of the j^th^ planting date (Planting Date 1 or Planting Date 2), respectively and the residuals were assumed to be independently and identically distributed, 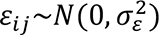. A Welch’s t-Test was used to compare grain fill per each inbred and storage interval combination to that of the control self-pollinations made on the same day. Finally, a Tukey *post-hoc* test was conducted across the three inbred lines and per combination of inbred and planting date to compare seed set between lines and over time per inbred line at an experiment-wise error rate (#x03B1;*_E_*) of 5%.

### 2.5 Experimental assessment of stored pollen across diverse inbred lines

To test the efficiency of the pollen collection method across inbred lines, pollen across 24 diverse inbreds among the major and sub-heterotic groups (White et al., 2020) was collected from the field and stored up to 24 hours prior to making pollinations. Pollen across each inbred line was collected when at least 50% of the plants for each line were shedding pollen. The pollen was then mixed with the PEEK-MP140 at a ratio of 1:5.

Pollinations were made the day pollen was collected (Day 0) and 24 hours after collection (Day 1). On each day, four LH244 plants with receptive silks were pollinated using the mixture and three self-pollinations of LH244 were made as controls. Images were acquired and used for visual rating and counting the number of kernels on each ear as described above. The effect of inbred on storage time was analyzed based on the average number of kernels over the four replicates using an ANOVA based on the model y_ij_ = Inbred_i_ + Storage_j_ + ε_ij_. Where Inbred corresponds to the effect of i^th^ line and Storage is the effect of the j^th^ storage interval. The residuals were independent and identically distributed, 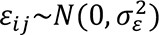. A two-sample Welch’s t-Test assuming unequal variance was used to compare grain fill per each inbred and storage interval combination to that of the control self-pollinations made on the same day. A Welch’s t-Test was also used to compare grain fill between days zero and one per inbred line.

## 3 RESULTS AND DISCUSSION

### 3.1 Assessment of storage substrate

The objective of this work was to develop and evaluate a method for cost-effective storage of maize pollen and efficient use of the pollen for breeding and genetics research. We observed that pollen that was stored without a substrate tended to quickly clump likely due to a chain-reaction of lysing pollen grains in contact with micro-environmental conditions. The literature also supports that mixing pollen with substrates could improve storability (Barnabas & Rajki, 1976). Two different substrates that supported successful pollen storage were initially identified, PEEK-MP140 and blue polyethylene microspheres (Figure 2). Of those substrates, the PEEK-MP140 was easily available and inexpensive and subsequently used for testing.

**Figure 2.**
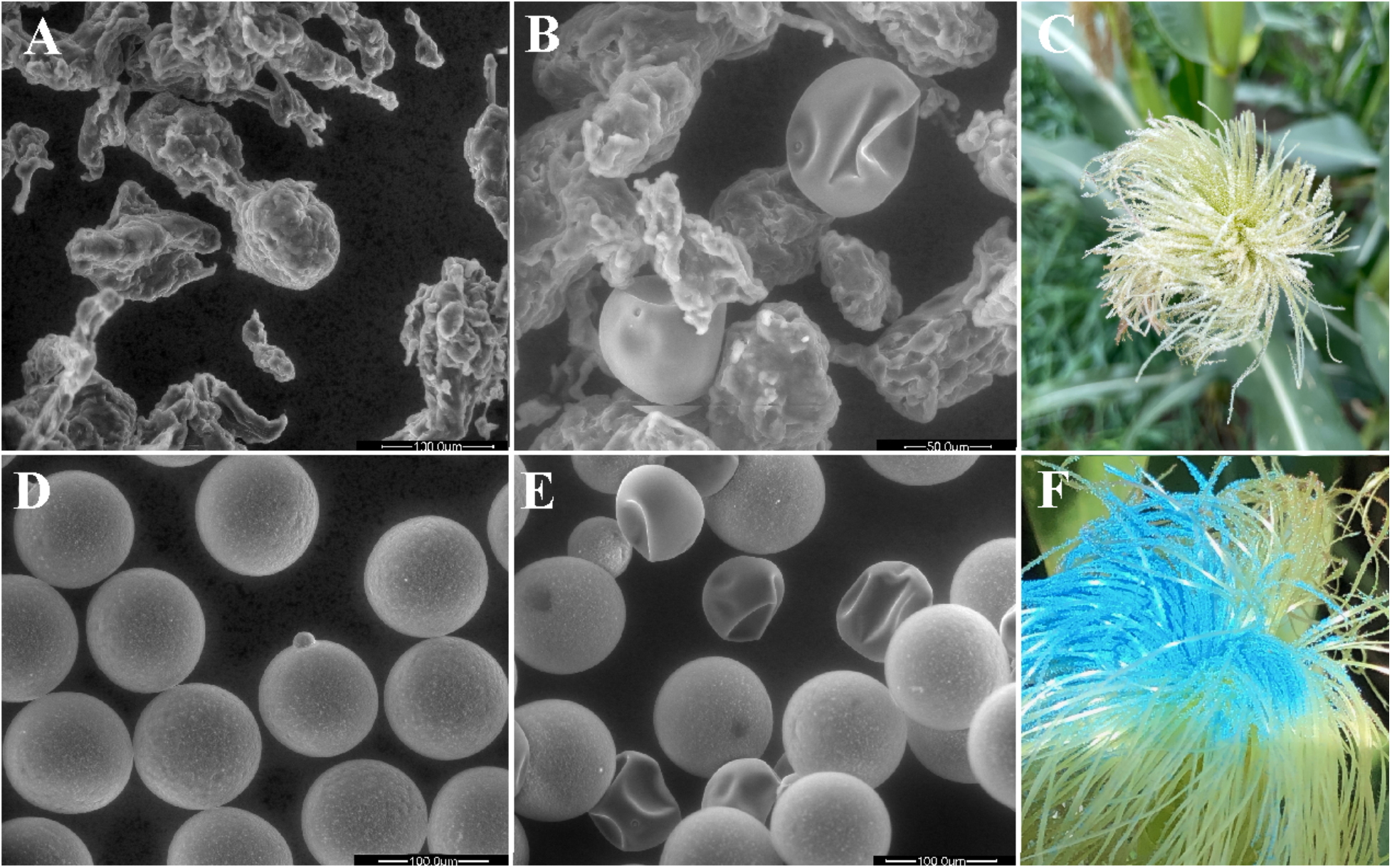
A) Scanning electron microscopy (SEM) images of a ground PEEK substrate called PEEK-MP140, B) SEM image of the PEEK-MP140 substrate mixed with pollen after 24 hours storage at approximately 6°C. D) SEM images of blue polyethylene microspheres (PEM) E) and SEM image of PEM mixed with pollen and stored for 24 hours. Example images of C) PEEK-MP140 and F) PEM mixed with pollen and applied to receptive silks after the mixture was stored for 24 hours at 6°C.

The hypothesis is that a substrate similar in size (approximately 90*μm to* 100*μm*) to typical pollen grains (Wodehouse, 1935; Jones & Newell, 1948) is more likely to form a homogenous mixture. If the substrate was larger than the pollen, the granules would sink to the bottom of the vessel and affect the homogeneity of the mix dispensed onto the silks of the seed parent. Based on this hypothesis, silica powders, polymer microspheres, diatomaceous earth, perlite powders and PEEK were initially evaluated for their ability to store maize pollen. Initial assessment demonstrated that PEEK-MP140 from PolyClean Technologies Inc. (Table 2) and blue microsphere polyethylene (PEM) could effectively store maize pollen (Supplemental Table S1).

**Table 1.** Inbred selection and heterotic group designation of 24 inbred lines

| **Inbred** | **Heterotic Group** |
| --- | --- |
| 3IIH6 | Iodent |
| 91BMA2SR* | B14 |
| FBLL | B73 |
| LH188 | Lancaster |
| LH198 | B73 |
| LH200 | B73 |
| LH223 | B14 |
| LH225 | B14 |
| NKH8431* | B73 |
| NP2011 | B73 |
| NP2031 | Flint |
| NP2151* | B73 |
| NP764 | B73 |
| NP942 | Iodent |
| PH06N | Iodent |
| PH09E | B37 |
| PH41E | Iodent Lancaster |
| PH44A | B37 |
| PHJ89 | Oh43 |
| PHN46 | Iodent Lancaster |
| PHR31 | Iodent |
| PHW03 | Flint |
| PHW20 | Flint |
| WQCD10 | B73 |

\*Inbred line was absent from White et al. (2020), so heterotic grouping was inferred based on pedigree information.

**Table 2.**
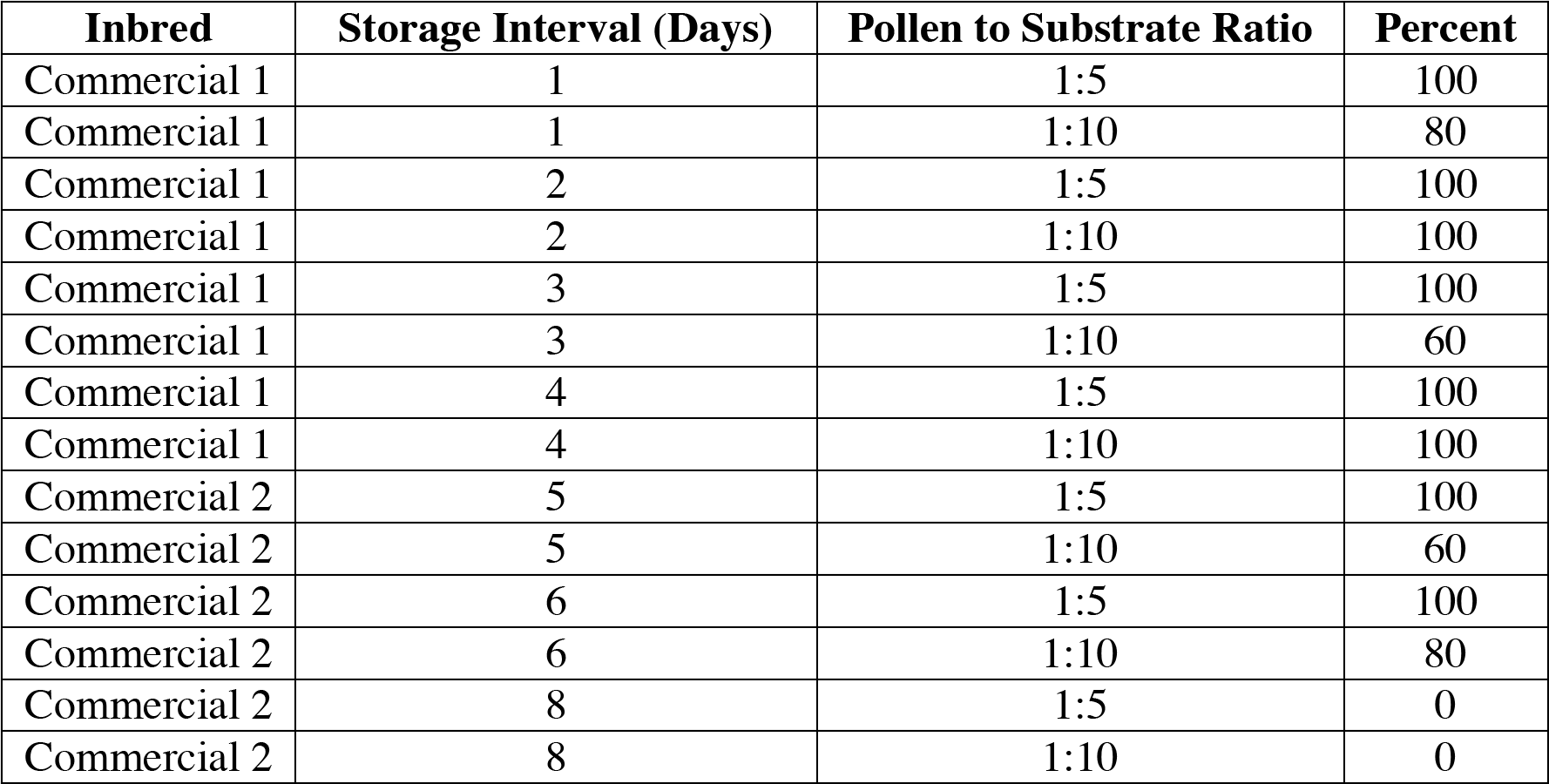
Percentage of ears out of five replicate pollinations with kernels at harvest that were pollinated with mixed pollen stored up to eight days using a 1:5 and 1:10 ratio of pollen to PEEK-MP140

Scanning electron microscopy allowed us to capture high-resolution close-up images of single pollen granules from the inbred line LH244 within each of the two substrates (Figure 2B and 2E). Observational analysis of the SEM images demonstrates that both substrates are similar in size to that of a single pollen granule but have distinct morphological characteristics (Figure 2). For example, the PEEK-MP140 substrate is approximately the same size as a single pollen granule, but each individual granule contains an irregular and non-consistent morphological shape (Figure 2A and 2B). Alternatively, each individual PEM particle is an identical sphere similar in size to a grain of pollen (Figure 2D and Figure 2E). In comparison, diatomaceous earth is a ground powder substantially smaller than an individual pollen grain. Diatomaceous earth and the silica powders failed to maintain pollen viable in initial tests (Supplemental Table S1). The PEEK substrate was acquired for approximately $0.07 per gram compared to $15.00 per gram for PEM, which was previously used for pollen cryopreservation (Barnabas & Rajki, 1976). Using a PEEK based product is a 214-fold decrease in cost compared to polyethylene substrates as used by Barnabas & Rajki (1976), improving the cost-effectiveness of this protocol for storing maize pollen.

### 3.2 Evaluation of maize pollen storability

The method for collection and storage of maize pollen was evaluated over three years beginning in 2020 using field experiments. Initial assessments of the method evaluated its utility for hybrid seed production in a breeding nursery and evaluated the effect of pollen concentration on seed set when the mixture was stored for up to eight days. Seed set was observed on two maize inbred lines pollinated after maize pollen was stored up to six days, but grain fill was not observed on day eight. A 1:5 ratio of pollen to PEEK-MP140 consistently generated more kernels per ear compared to a 1:10 ratio and grain fill decreased over storage time (Table 2). After the pollen mixture was stored for six days, only a few kernels were detected and just scattered throughout the ear (Supplemental Figure S2B and S2D). Overall, our results demonstrated that a sufficient proportion of maize pollen granules are viable up to six days of storage if quickly mixed with PEEK-MP140 as grain fill was observed on approximately 50% of the ear (Supplemental Figure S2). When the pollen mix was stored beyond 24 hours, a greater concentration of pollen to medium increased the number of kernels produced (Table 2), suggesting that the ratio of pollen to PEEK-MP140 is a critical variable in the procedure and pollen concentration influences seed set.

The experimental results from the summer of 2020 demonstrated that the method for pollen collection and storage can generate hybrid seed after six days of storage. However, we found that the pollen concentration can influence seed set. With this information, we implemented the procedure for seed production in our maize breeding and genetics research program beginning in 2021 and consistently observed ears with complete grain fill at harvest (Supplemental Figure S3). That same summer we harvested approximately 1.2 million kernels across 1,506 nursery rows when six to fifteen ears per row on average were pollinated with mixed pollen. We have observed that collecting and mixing pollen in a storage substrate increases the number of seed parents that can be pollinated compared to traditional hand crossing. From routine utilization of this method within our breeding program, five milliliters of pollen collected from five to 25 tassels, dependent on inbred, can produce a pollen mix that can pollinate more than 200 plants or more than 10 pollinations per tassel.

### 3.3 Evaluation of timing of pollination

In 2021, the importance of the concentration of pollen to PEEK-MP140 was evaluated. Pollen from the inbred line PHP02 was collected and mixed with PEEK-MP140 using both a 1:5 and 1:10 dilution. The two different mixtures were stored for 48 hours and then each mixture was applied to ears of PHP02 plants with receptive silks. The pollen was applied every hour between 7:00 A.M. and 12:00 P.M. and we observed that the timing of application did not significantly influence percent grain fill (P-value > 0.05) while the ratio of pollen to substrate did significantly influence grain fill when the mixture was stored for 48 hours (P-value < 0.05) across these times and days of storage (Supplemental Table S2). On average, using a 1:5 ratio mixture led to a larger number of ovules successfully pollinated based on visually rating of percent grain fill relative to a 1:10 ratio (Figure 3). Having more pollen in the mix may be adventitious as having more granules in the mixture increases the probability that a viable pollen granule will land on a silk, germinate, and fertilize an ovule (Heslop-Harrison, 1979). A significant difference between the two ratios was only observed when pollinations were made at 9:00 A.M. or 10:00 A.M. and that difference changed depending on if the mixture was stored for 24 or 48 hours (Figure 3).

**Figure 3.**
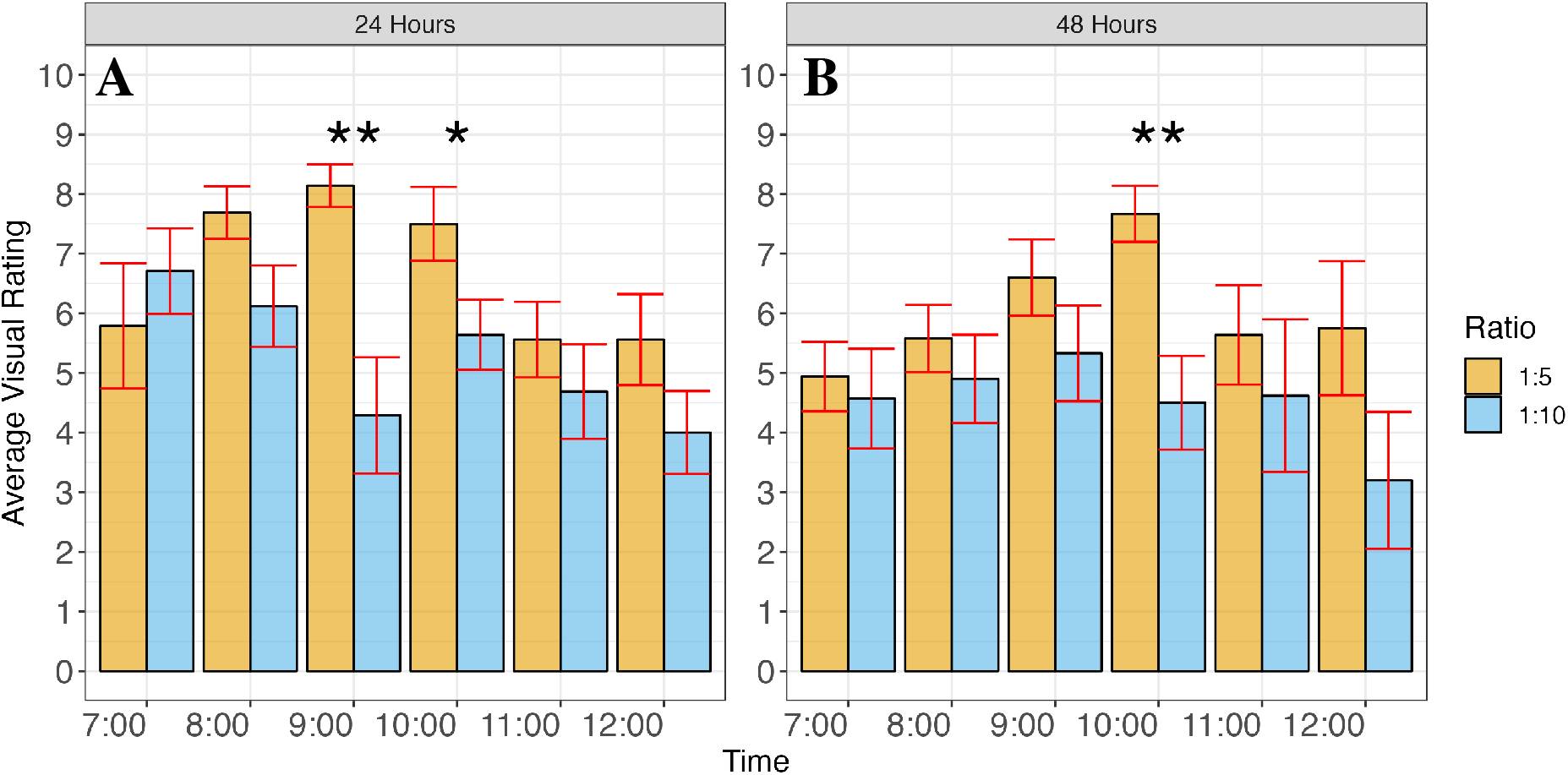
Grain fill at harvest based on visual rating for percent grain fill for the evaluation of PHP02 pollen mixed and stored in PEEK-MP140 for A) 24 and B) 48 hours prior to being applied to PHP02 plants with receptive silks. Orange bars show the 1:5 ratio of pollen to PEEK-MP140 and blue bars show the 1:10 ratio with red standard error bars. ‘*’ and ‘**’ correspond to P-values < 0.05 and P-values < 0.01 respectively based on a Welch’s t-Test between the 1:5 and 1:10 ratio per time and hours of mixed pollen storage.

The proportion of the ear with grain after the pollen mixture was stored for 24 hours was maximized when pollinations were made during the mid-morning or between 9:00 A.M. and 10:00 A.M. (Figure 3A). However, we generally observed that the average grain fill between each combination of pollen concentration and timing of application per storage interval was not significantly different based on a 5% experimental wise error rate using a Tukey *post-hoc* analysis. Storing pollen from inbred line PHP02 resulted in a decrease in grain fill between days one and two but even after 48 hours of storage, grain was observed on over 50% of the ear (Figure 3). Practically, seed generation via hand crossing where pollen in the tassel bag is carried to the seed parent would not be possible most days prior to late morning or early afternoon within our geographic region as heavy moisture in the bag of pollen would lead to pollen bursting and dehiscence of new pollen would not yet have occurred due to insufficient heat (Bair & Loomis, 1941). Heavy rainstorms can also lead to total saturation and loss of the tassel bag, prolonging the period from silk emergence to pollination, potentially leading to a loss in grain fill due to reduced silk receptivity associated with aging of the flower (DuPlessis & Dijkuis, 1967; Wych, 1988; Bassetti & Westgate, 1993). Using stored maize pollen for crossing in a breeding program has the potential to mitigate these issues by allowing pollen to be collected from plants grown in a controlled environment or from a previous day and transported to a field when the silks on the ear are at prime receptivity.

### 3.4 Evaluation of pollen storability across planting dates

In 2022, our method was directly compared to the current standard self-pollination procedure as a control. Pollen from the inbred lines LH244, PH24E, and LH287 was collected and stored then used to pollinate LH244 plants with receptive silks. The number of kernels harvested from the controls across two planting dates was used as a baseline to compare relative grain fill success. Among the controls, seed production was lower for the first planting compared to the second planting as the average number of kernels observed on the ears of the controls was 250 and 157 kernels per ear for the first and second planting date, respectively (Figure 4). As a percentage of the control, seed set using collected and stored pollen was lower on average for the first planting but outperformed the controls on day zero and one for the second planting (Figure 5).

**Figure 4.**
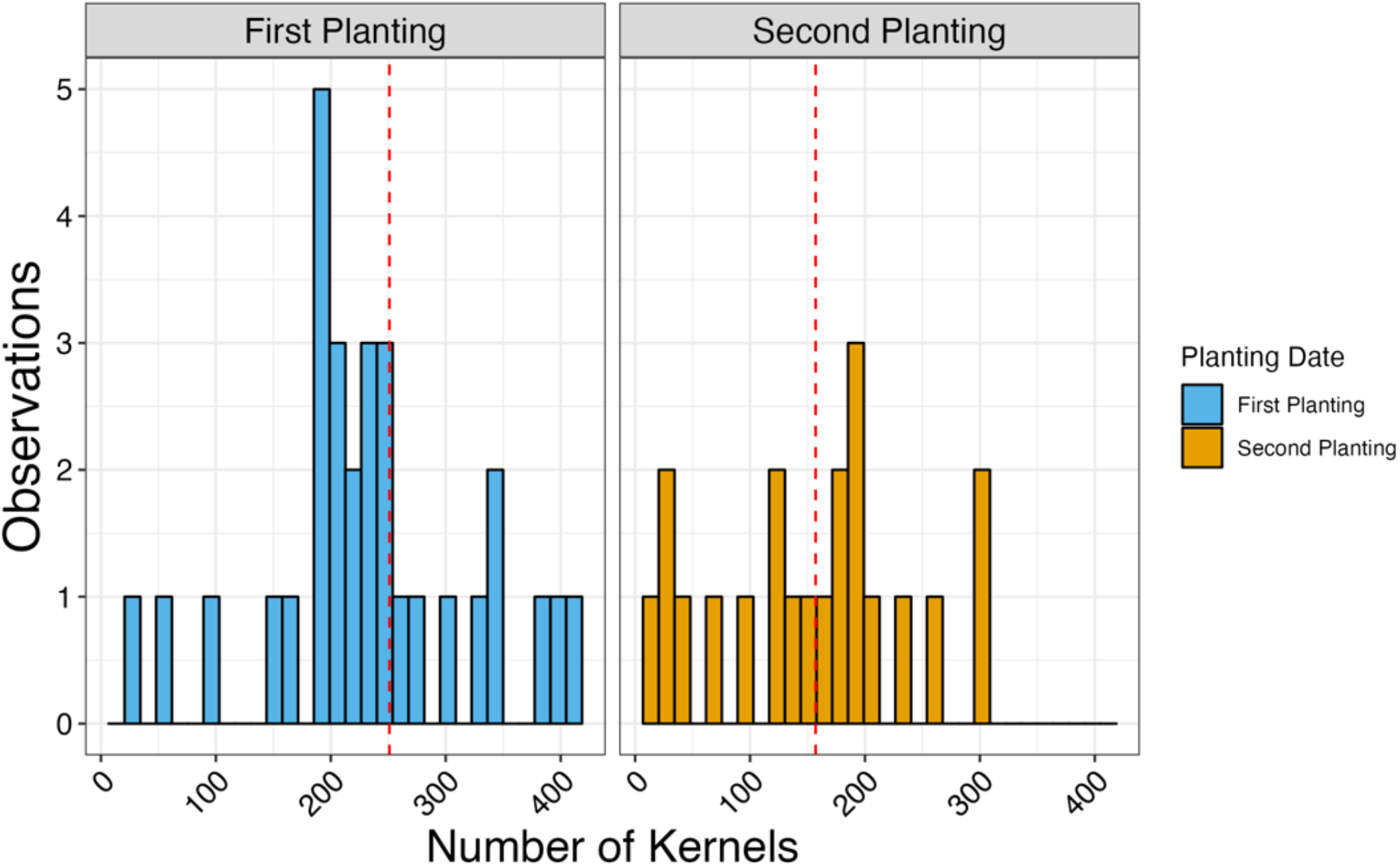
The average number of kernels harvested across planting dates for the controls shown by the dashed red line. Controls are defined as the self-pollination of the inbred line LH244.

**Figure 5.**
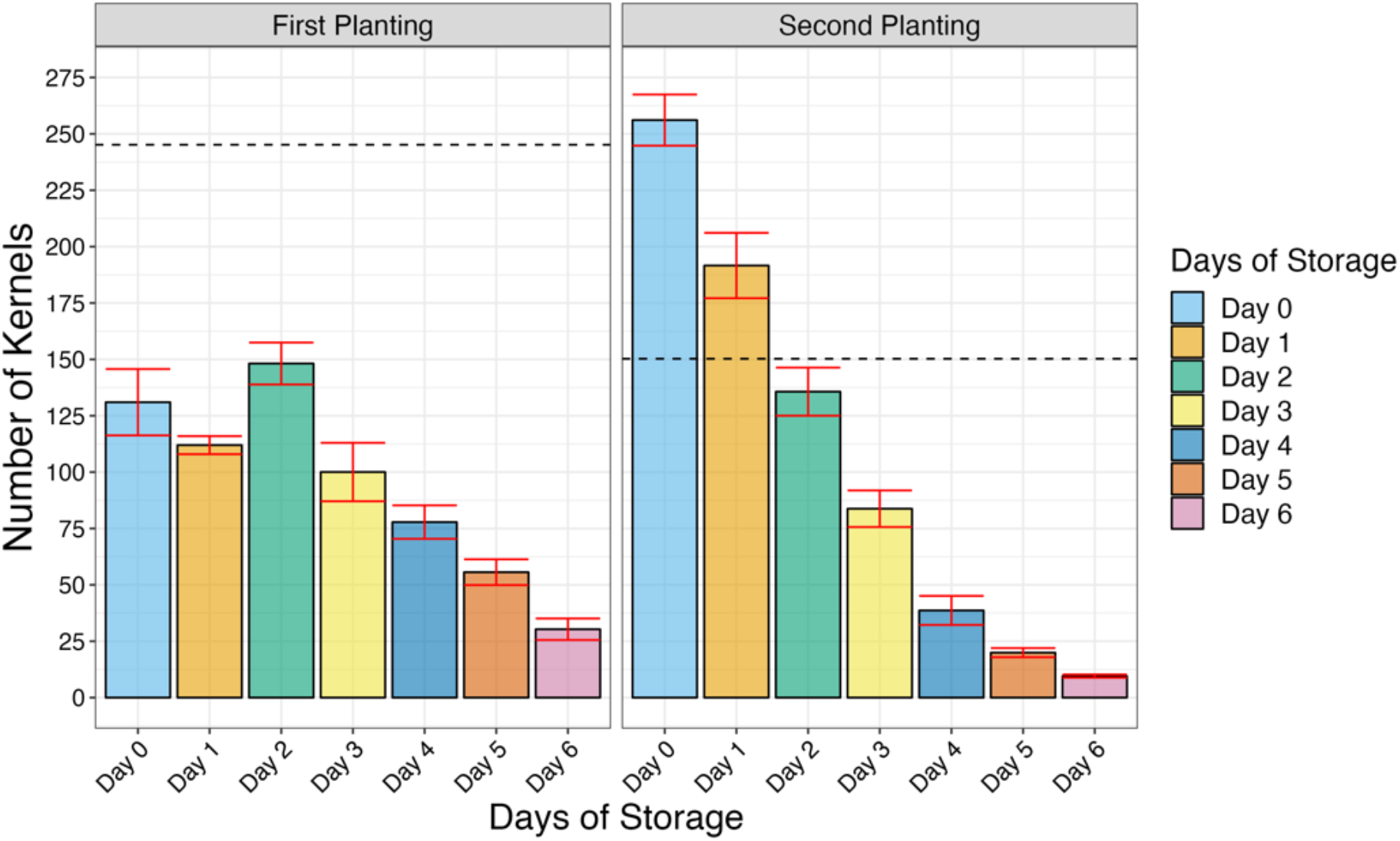
The average number of kernels harvested among ears pollinated with mixed pollen stored up to six days using a 1:5 ratio compared to the average number of kernels harvested from the controls per planting date. The average number of kernels harvested for the controls per planting date is shown by the horizontal black dashed line. Results shown for two different planting dates that correspond to an early (First Planting) and late planting (Second Planting) within Central, WI. Red bars show the standard error of the mean across the three inbred lines per storage interval. Bars are color coded by days of storage.

Grain fill appeared to dramatically decrease between day five and six (Figure 5) and a maximum 34 kernels on average were harvested after five days of storage, so the average number of kernels per ear across the six replicate pollinations analyzed only considered days zero to five of storage. Additionally, no pollinations were made after six days of storage for the second planting as no silks were available due to high Corn Rootworm Beetle (genus *Dabrotica*) pressure.

Storage interval was significant while planting date did not significantly affect grain fill (Table 3). In general, more kernels were harvested from the standard controlled pollinations compared to the ears pollinated with the mixed pollen after maize pollen was stored for 48 hours (Figure 5). However, the mixed pollen method was highly effective and there were multiple examples where the mixed pollen outperformed the control. For example, more kernels were harvested from ears pollinated with stored PH24E pollen than the controls for the second planting on days zero to three. For LH244, a greater number of kernels were harvested using mixed pollen compared to the control pollinations on day one and two for the first planting and on day zero and one for the second planting. These results suggest that the method has the potential to outperform the traditional self-pollination procedure even when maize pollen is stored up to 72 hours.

**Table 3.** ANOVA for the average number of kernels per ear and visually rated percent grain fill per ear when pollen from the inbred lines LH244, PH24E, and LH287 is collected and stored for five days and used to pollinate the inbred line LH244

|  | Number of Kernels |  | Percent Grain Fill |  |
| --- | --- | --- | --- | --- |
|  | F | P-value | F | P-value |
| Days of Storage | 3.519 | 0.001 | 4.160 | 0.006 |
| Planting Date | 0.418 | 0.523 | 0.172 | 0.680 |

The experimental results in 2022 demonstrated that at least 50 kernels can be harvested after mixed pollen is stored for five days (Figure 5). For the second planting date, pollen from PH24E successfully generated at least 50 kernels after five days of storage (Supplemental Table S3). Variation in seed set among ears pollinated with the three different inbred parents is likely due to technical variation introduced by a day effect and differences in timing of collection. For example, LH244 is later maturing compared to PH24E and LH287, so pollen was collected two days after the latter two inbred lines for the first planting date. For the second planting date, LH287 and LH244 was collected in the late afternoon while PH24E pollen was collected the following day at mid-morning. However, after five days of storage, 100 kernels or approximately 50% of the ear was covered with grain at five days of storage when plants with receptive silks were pollinated with PH24E pollen, and 51 kernels were still harvested when plants were pollinated with LH244 pollen. When plants were pollinated with pollen from LH244 or PH24E, almost 100 kernels were harvested after four days of storage for the first planting date (Supplemental Table S3). These results suggest that efficiently mixing the PEEK-MP140 substrate with pollen adjacent to the field and quickly transporting the mixture to a cool environment at approximately 6°C has the potential to maintain enough pollen granules viable for sufficient seed production in a breeding and genetics program.

Interestingly, when we averaged across the three pollen parents for this analysis, we observed a greater number of kernels harvested using mixed pollen compared to the controls on days zero and one for the second planting date. These results suggest that the method can work effectively for collecting pollen even late in the growing season within our geographic region. High temperatures are known to accelerate the rate of pollen desiccation via rapid pollen-water loss and there is a negative correlation between pollen desiccation rate and temperature (Schoper et al., 1987a and 1987b; Roy et al., 1995). Given this biological understanding and our experience using the method for seed production in our breeding program, we recommend collecting pollen for storage in the morning when the tassel bag is dry and right at the start of dehiscence to maximize pollen quality for storage and use.

Additional external environmental factors such as high insect pressure caused by Corn Root Worm beetles could have contributed to both the plant-to-plant variation in grain fill for a given storage treatment and potentially introduce contamination. Insect pressure was substantial in the second planted material in 2022. Plant-to-plant variation can have a large effect on overall seed set due to differences in silk brush receptivity between ears (Westgate & Boyer, 1986; Aylor, 2004). The controls exhibited variation in grain fill both within and between planting dates (Figure 4) suggesting that factors outside of the methods described for collection and storage of maize pollen influence the number of kernels harvested during seed production. Therefore, the described method appears effective for seed production throughout the growing season and is not limited by planting date.

The variation in grain fill between control plants and plants pollinated with mixed pollen was similar up to four days of storage for both plantings. By day five for the first planting, the standard deviation in grain fill was greater among the controls compared to the pollinations made using the mixed pollen. Interestingly, the variation in grain fill can potentially be reduced using stored pollen compared to self-pollinations as exemplified on day one for the second planting, where the average grain fill standard deviation when using stored pollen was 60.10 kernels compared to 67.30 kernels for the controls. These results suggest that using stored pollen may help reduce plant-to-plant variability in grain fill during seed production.

Collecting maize pollen directly from tassels, mixing the pollen with a substrate, and directly using the mixture to pollinate ears with receptive silks has the potential to generate grain fill similar that if mixed pollen was stored for five days (Table 4). For example, while grain fill was lower at day five relative to day zero when pollen was collected from inbred line PH24E, the number of kernels on the ear between those two days was not significantly different (Table 4). These results were also supported by our binary assessments of grain fill in 2020 where on average, 50% of the ear exhibited grain fill at both day zero and five (Table 2). Jones and Newell (1948) observed that seed set dramatically decreased after two days when the inflorescence containing unreleased pollen was refrigerated. However, the method that we describe allows storage for at least five days. Additionally, over 25% of the ear can still be filled with grain after six days and up to 20 kernels were observed on the ear after eight days of storage (Supplemental Table S3) with the amount of pollen mixture applied.

**Table 4.**
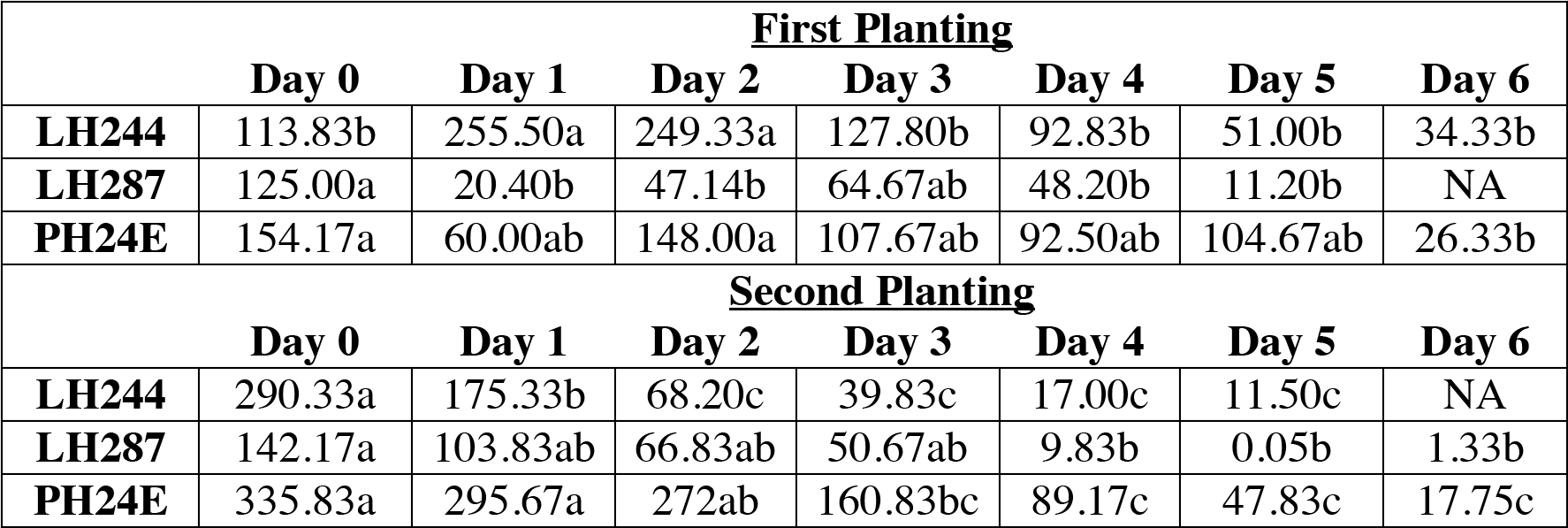
Average grain fill over time per inbred line and planting date based on the number of kernels per ear when mixed maize pollen is stored out to six days. Letters represent significant differences in grain fill between day zero and six for each inbred line per planting date based on Tukey Honest Significant Difference (HSD) test at a 5% experiment-wise error rate. Missing values (NA) represent days where no pollinations were made due to inclement weather.

Future investigations using pollen germination assays could help estimate the proportion of viable granules in the mixture at five days or greater of storage to determine if a greater concentration of pollen to media is required for storage beyond five days. Additionally, an initial pollen germination assay could help determine if the variability in grain fill over time (Table 4) is associated with the number of viable pollen granules harvested during collection. However, the goal of this paper was to describe a method for collecting maize pollen and demonstrate the utility of using stored maize pollen for seed production in breeding programs, so the aforementioned two hypotheses are a subject of future research.

Seed set was observed on LH244 ears pollinated with maize pollen collected after nine days of storage with a maximum of 12 kernels per ear observed on day nine (Supplemental Table S3). Therefore, the procedure can lead to seed production with some level of success after nine days of storage. These results are consistent with the findings of Jones and Newell (1948) who also observed seed set on maize cultivars pollinated with pollen stored for nine days. In comparison to the work of Jones and Newell (1948), our procedure works by diluting the concentration of pollen via mixing the pollen with a PEEK based substrate to increase the number of plants that can be pollinated per bag of collected pollen.

### 3.5 Evaluation of pollination effectiveness across diverse maize inbreds

To further explore genetic differences in pollen storability, we evaluated the utility of our process across 24 inbred lines (Table 1) that represented a wide variability within US Dent germplasm (White et al., 2020). Impe et al. (2020) observed that *in vitro* pollen germination between inbred lines PH207, A188, B73, and A183 ranged from 0.2% to 4.5%, suggesting that genetics could influence storability. However, our method worked effectively across all 24 inbred lines with an average of 67 to 245 kernels per ear harvested at day zero and a maximum of 103 to over 300 kernels harvested per ear (Table 5). Grain fill decreased on average between day zero and one from 158 (±10.46) to 121 (±16.01) kernels per ear. However, this is still equivalent to observing approximately 45% grain fill per ear on average across the 24 inbred lines.

**Table 5.**
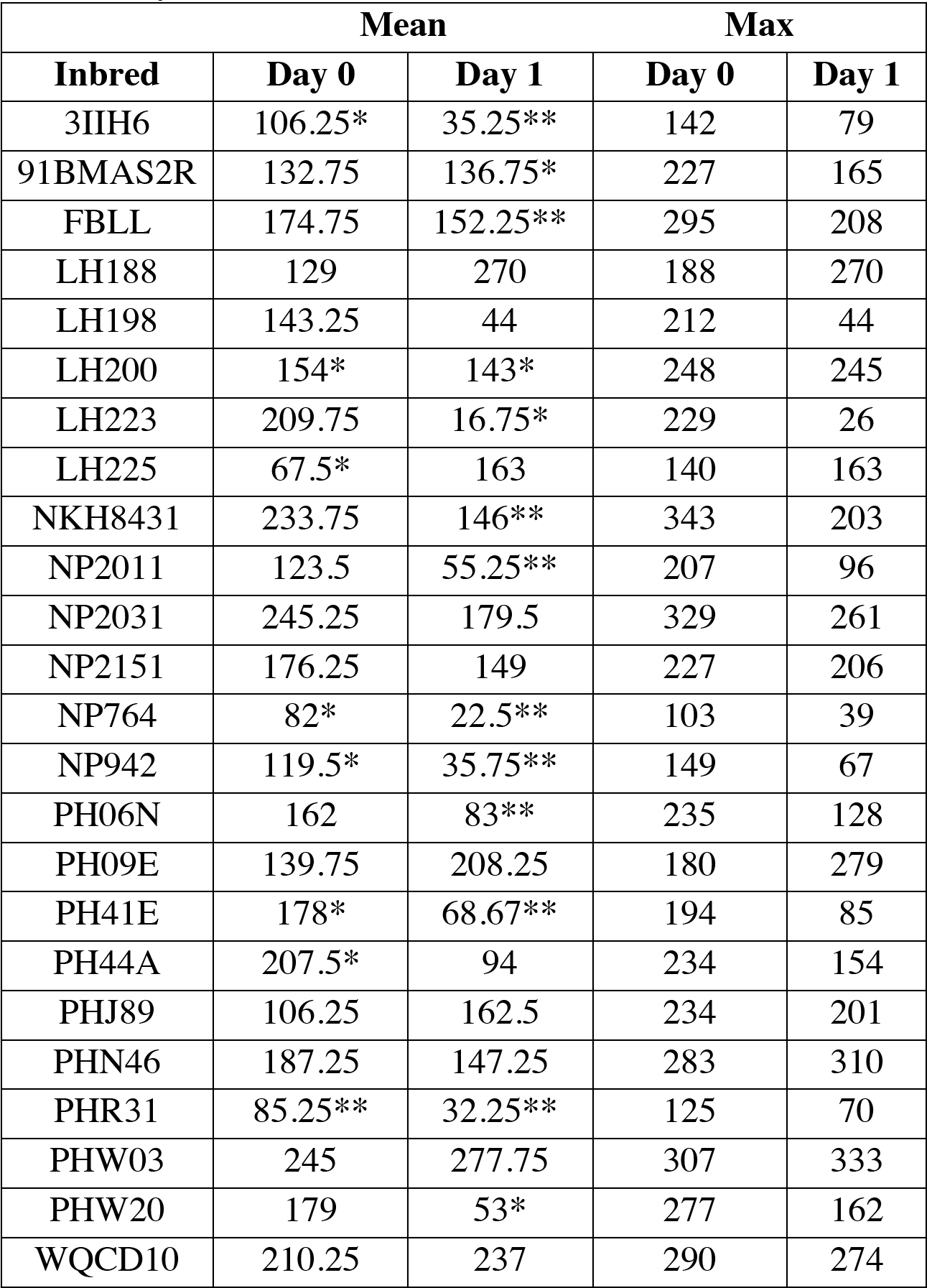
Average mean and maximum grain fill over four replicates when maize pollen was collected and mixed with PEEK-MP140 across 24 different inbred lines and used to pollinate LH244 silks immediately (Day 0) or after mixed pollen was stored for 24 hours (Day 1). ‘*’ and ‘**’ correspond to P-values < 0.05 and P-values < 0.01 respectively based on Welch’s t-Test comparing mean grain fill among ears pollinated with mixed pollen to the control self-pollinations on the same day.Table 6. ANOVA describing the effect of inbred line and days of storage on average grain fill when pollen was collected from 24 different inbred lines and immediately used to pollinate LH244 or stored for 24 hours prior to pollination

On average, grain fill decreased between days zero and one as expected and significantly affected grain fill at harvest (Table 6). Seed set per storage interval was equivalent to the controls for 66% of the inbred lines on day zero and equal to the controls among 45% of the inbred lines on day one (Table 5). These results demonstrate that the efficient procedure for collection and storage of maize pollen works effectively across diverse genetic backgrounds within the US Dent germplasm.

We tested if the inbred line used as a pollen parent had a significant effect on grain fill using an analysis of variance. Inbred line did not have a significant effect on seed set (Table 6) and suggests the method is not limited by the choice of inbred line. Although, we did observe some variation in grain fill across the 24 inbred lines used as pollen parents, much of the variation could be the result of a day effect as pollen was collected across four different days due to variation in days to anthesis between the inbred lines (White et al., 2020).

**Table 6.**
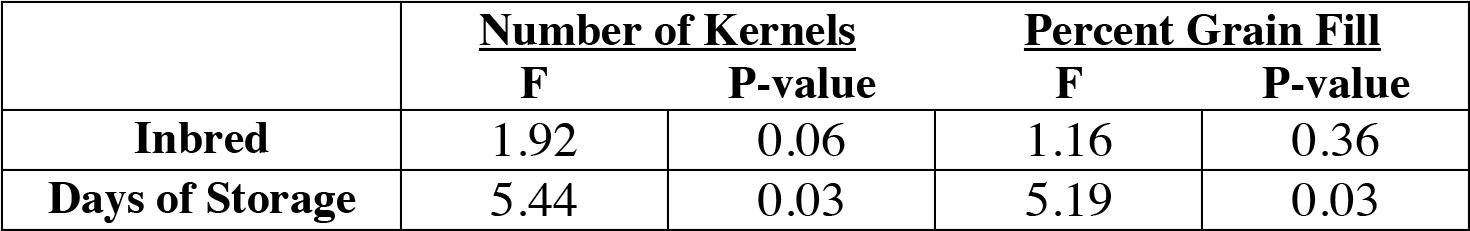
ANOVA describing the effect of inbred line and days of storage on average grain fill when pollen was collected from 24 different inbred lines and immediately used to pollinate LH244 or stored for 24 hours prior to pollination

Daily differences in humidity and temperature across the four collection dates could have influenced pollen desiccation during collection (Schoper et al., 1987a and 1987b; Roy et al., 1995). Additionally, differences in the water content of the silks among LH244 plants used as the seed parent could have reduced receptivity (Bassetti & Westgate, 1993) and led to variation in grain fill when plants are pollinated using the diverse set of inbred lines in this experiment. From routine utilization within our maize breeding program, we have not observed any limitations in the method due to the choice of inbred led line during hybrid seed generation. As an example, in one seed production nursery in Verona, WI in 2022, we used this method to collect pollen across 30 unique inbred lines that included both ex-PVPs, current commercial inbred lines, and publicly developed double haploids from the WI-SS-MAGIC population (Michel et al., 2022). Collecting pollen across this diverse germplasm led to the generation of over 230 hybrids when mixed pollen was directly applied to plants with receptive silks or stored for 24 hours prior to application.

### 3.6 Conclusion

The purpose of the current study was to develop and evaluate a practical method for cost-effective and efficient collection and storage of maize pollen. A substrate was identified, PEEK-MP140, which is approximately the size of an individual pollen granule and is useful to produce a homogenized suspension that supports extension of pollen viability. Even after six days of storage, the method has the potential to maintain enough viable pollen granules such that at least 50 maize kernels can be harvested per ear on average (Supplemental Table S3) and this method works across a diverse set of maize inbred lines (Table 5). While maize pollen can be maintained in a polyethylene-based substrate and kept in liquid nitrogen for later use (Barnabas & Rajki, 1976; Barnabas et al., 1988), this method is expensive and lacks efficiency. The method demonstrated here mitigates the latter two issues by utilizing a PEEK based media without deep-freezing. Using this method, stored maize pollen could be routinely utilized for seed production in a breeding program or genetics research.

Storage of maize pollen would facilitate crossing of germplasm with maturity differences that complicate regular planting and crossing. In these cases, maize pollen from an early-flowering inbred line could be collected, mixed, stored, and applied to silks of the late-flowering parent at the time they are receptive. Planting ‘delayed rows’, or additional rows sometime after the initial planting to increase the probability of synchronous pollen shed and silk emergence, is a widely used practice but has logistical complexities and is not always a viable strategy when new germplasm with unknown flowering characteristics is being used.

Efficient seed production is vital for plant breeding and genetics programs but is labor intensive and expensive. However, the time and cost of seed production can be reduced by collecting pollen and storing it in an appropriate substrate at a reduced concentration as it is estimated that over a million pollen grains are produced in a single tassel, but only 200 to 300 viable granules are needed to fertilize all the ovules on an inbred line. The idea of using stored maize pollen in the context of a breeding program has been explored since the early 1920s (Knowlton, 1922) but has had limited utility due to the cost, complexity, and repeatability of the process. We demonstrate a simple and cost-effective process that has practical utility for routine seed generation in breeding programs and genetics research.

## ACKNOWLEDGEMENTS

The authors acknowledge Tim Osterhaus for assistance with data collection and harvest of the Summer 2022 experiments, Dr. Jose Varela for helpful discussions and recommendations during image acquisition of the 2021 samples, and the USDA Germplasm repository for providing useful germplasm. The work was supported by the National Institute of Food and Agriculture (NIFA), United States Department of Agriculture (USDA), and USDA Hatch Projects 1022702 and 1015851. In addition, this research was partially supported by the National Institute of Food and Agriculture and United States Department of Agriculture Hatch grants 1020442 and 1013262.

## SUPPLEMENTAL MATERIAL

Supplemental File S1 provides a step-by-step list of instructions for executing the described method of pollen collection, storage, and application. Supplemental Figure S1 shows the visual rating scale used for determining percent grain fill. Supplemental Figure S2 provides example of ears at harvest pollinated with mixed pollen for the evaluation of maize pollen storability experiment. Supplemental Figure S3 provide examples of harvested ears pollinated with mixed pollen during routine utilization for hybrid seed production within our maize breeding program. Supplemental Table S1 has information on the number of kernels harvested during the assessment of storage substrate experiment. Supplemental Table S2 has the ANOVA table for the evaluation of timing of pollination experiment. Supplemental Table S3 provides the summary statistics per each inbred line, storage interval, and planting date for the evaluation of pollen storability across planting dates experiment.

## Abbreviations

ANOVA: analysis of variance
DAP: days after pollination
PEEK: polyetheretherketone
PEM: polyethylene microspheres

