## Supplemental File S1 for "A practical method to improve the efficiency of pollination in maize breeding and genetics research"

### SUPPLEMENTAL METHODS

#### Step 1 - Pollen Collection

1. Place a tassel bag on the florescence of inbred lines shedding pollen 24 hours prior to collecting, mixing, and storing pollen.
    a. The quantity of pollen collected varies by inbred and is affected by the environment. However, we have consistently observed that one tassel can pollinate 10 plants with receptive silks using mixed pollen.
2. After 24 hours and when the temperature is appropriate for pollen dehiscence, place your hand at the base of tassel bag and carefully bend over the plant such that the tassel is approximately parallel to the ground.
    a. Tap the tassel three times with your hand to shake off all fresh pollen.
    b. Carefully remove from the tassel bag. During removal, shake the tassel gently back and forth to maximize the amount of pollen collected.
        i. Quickly, completely close the bag following removal. The bag can be closed by folding over the top part of the bag approximately 1.5” twice (Figure 1A).
3. If possible, transport the bags with fresh pollen to a covered and shaded environment adjacent to the field to reduce the rate of pollen desiccation.
    a. Currently, work efficiently but carefully to avoid pollen desiccation while still minimizing contamination.
4. For each bag with pollen, dump the pollen first through a stainless-steel strainer to remove anthers and large debris.

   a. A common kitchen strainer works sufficiently for this step. Our breeding program commonly uses a Mainstays 3" Mini Stainless Steel Fine Mesh Strainer Basket for this procedure.
      i. For removing pollen from the tassel bag, place the strainer on top of a plastic container to catch the pollen (Figure 1B). Any plastic container that is free of contamination will work sufficiently.
5. Take the sieved pollen and gently pour it through a fine mesh.
   a. Our breeding program commonly uses a Tansoole Experimental Sieve of size 100 mesh (0.154 mm) (Figure 1C) and have also observed a size 80 mesh ( 0.180 mm) to work. In general, the size of the mesh should allow fresh non-clumped pollen to flow through while capturing clumped pollen and small debris.
   b. Gently tap the sides of the sieve to help the pollen go through the mesh. The goal of this step is to remove clumped pulled or any other debris missed during the initial sieve.
      i. Throw out any clumped pollen that did not go through the mesh and proceed to Step 2.

**Step 2 - Pollen Mixture**

1. The sieved pollen is then mixed with the medium at an appropriate concentration.
   a. Measure out the quantity of sieved pollen using a 50 mL centrifuge tube. Generally, a funnel is used at this step to help pour the pollen from the sieve into the 50 mL centrifuge tube.

        i. Make a note of the quantity of fresh sieved pollen in the tube and dump the pollen into a 120 mL (4 oz) Sterile Specimen Container.
    b. Based on the quantity of freshly collected pollen, measure out the appropriate quantity of PEEK-MP140 substrate.
        i. Our maize breeding and genetics program has used a ratio of one to five (1:5) pollen to PEEK-MP140 (Figure 1D) during routine use of collected and mixed pollen for seed production.
2. Gently dump the medium into the specimen container with the pollen and place the lid onto the container.
    a. Confirm that the lid is sealed tightly before proceeding.
    b. Hold the container horizontally and gently rotate the container approximately five times or until a homogenized mixture is created (Figure 1E).
    c. If no pollinations are to be conducted on the day of collection, directly place the mixture in a cold room or refrigerator that is between approximately 4°C and 6°C. Throughout our field experiments, storing mixed pollen at 6°C in a walk-in cold room worked sufficiently.

**Step 3 - Application**

1. Only ear shoots previously covered prior to silk emergence or away from any field shedding pollen should be used to avoid contamination.
2. If the mixture was stored, gently rotated it three times prior to aliquoting to the application vessel to ensure a homogenized mix.
3. Then, place the freshly mixed or stored pollen mixture into an application vessel.

a. For pouring the mixed pollen, place a funnel in the application vessel and dump in the mixture.

b. Our breeding program commonly uses 2.7 oz glass spice container with approximately 13 one-millimeter sized diameter holes (Figure 1F). However, additional application vessels with a similar size and number of holes for application pollen will work sufficiently.

    i. For example, we have observed that a 50 mL centrifuge tube with roughly 10 one-millimeter diameter holes works sufficiently.

c. Now, the mixture can be applied to receptive silks in the field or green house. When the mixed pollen is removed from the refrigerated environment, keep the mixture in a cool environment, if possible, to prevent desiccation and quickly as possible begin to pollinate targeted receptive silks.

    i. For example, placing the application vessel with mixed pollen in a cooler that was also kept in a cool environment may help reduce the rate of pollen desiccation.

        1. Ensure that no water can enter the application vessel during transportation to the target plants.

d. Apply the mixture to silks by delivering three ‘shakes’ of mixed pollen from the vessel to each ear where a ‘shake’ is defined as the movement of the applicators arm from a 90° to 45° angle when the vessel is maintained perpendicular to the forearm (Figure 1G).

    i. A greater quantity of mixed pollen may be adventitious if the silks are older, or the mixed pollen is stored longer. However, three complete 'shakes' from the application vessel should completely cover all receptive silks (Figure 1C).

e. Place a tassel bag immediately over the pollinated ear shoot and staple together the bag on the opposite side to prevent pollen from adjacent plants landing on the pollinated silks (Figure 1H).
    i. Continue the application process until all plants of interest with receptive silks are pollinated.
f. In Figure 1I are example of maize ears at harvest that were pollinated with mixed pollen in comparison to a representative control self-pollination (far left ear). Pollen was collected and then directly used to pollinate receptive silks (second ear from left) or stored out to four days before being used to make pollinations.
