## Supplemental Figure S1 to S3 and Supplemental Table S1 to S3 for "A practical method to improve the efficiency of pollination in maize breeding and genetics research"

**Supplemental Figure S1.** A visual rating scale from one to ten used to measure percent grain fill per ear.

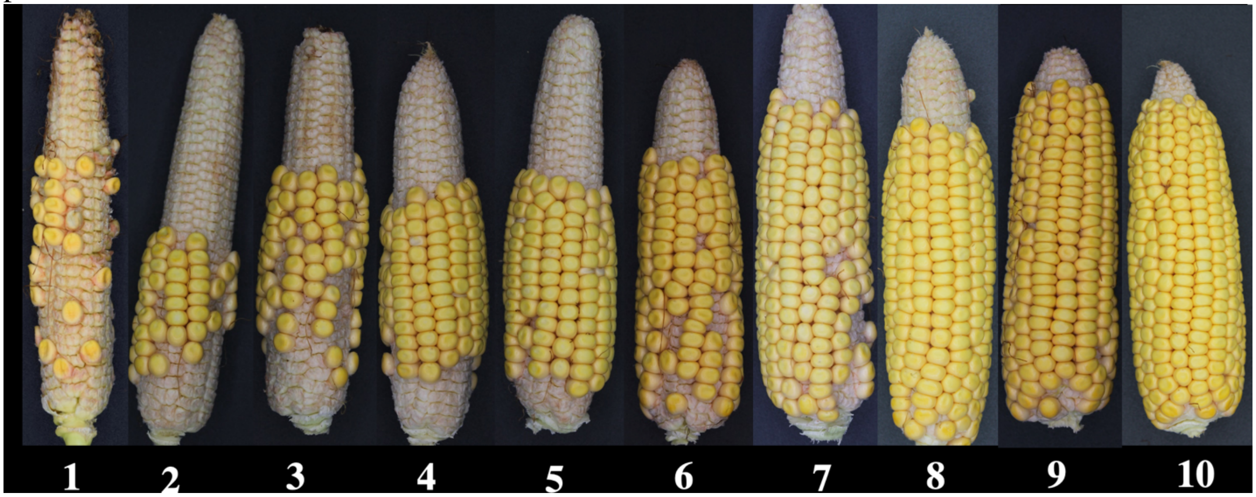

**Supplemental Figure S2:** Maize ears collected and harvested in 2020 that were pollinated with pollen from a line heterozygous for purple pigmented kernels mixed with PEEK-MP140 at a ratio of 1:5 and 1:10 pollen to substrate and stored between 24 hours and eight days. A) and B) show the first side of one representative maize ear that was pollinated and C) and D) show the same ears rotated 180°. In the image labels per board, the first number represents the number of days of storage and the second number presents the pollen-substrate ratio as 1:5 or 1:10. All ears were pollinated using a pollen mixture from the same 50 ml tube.

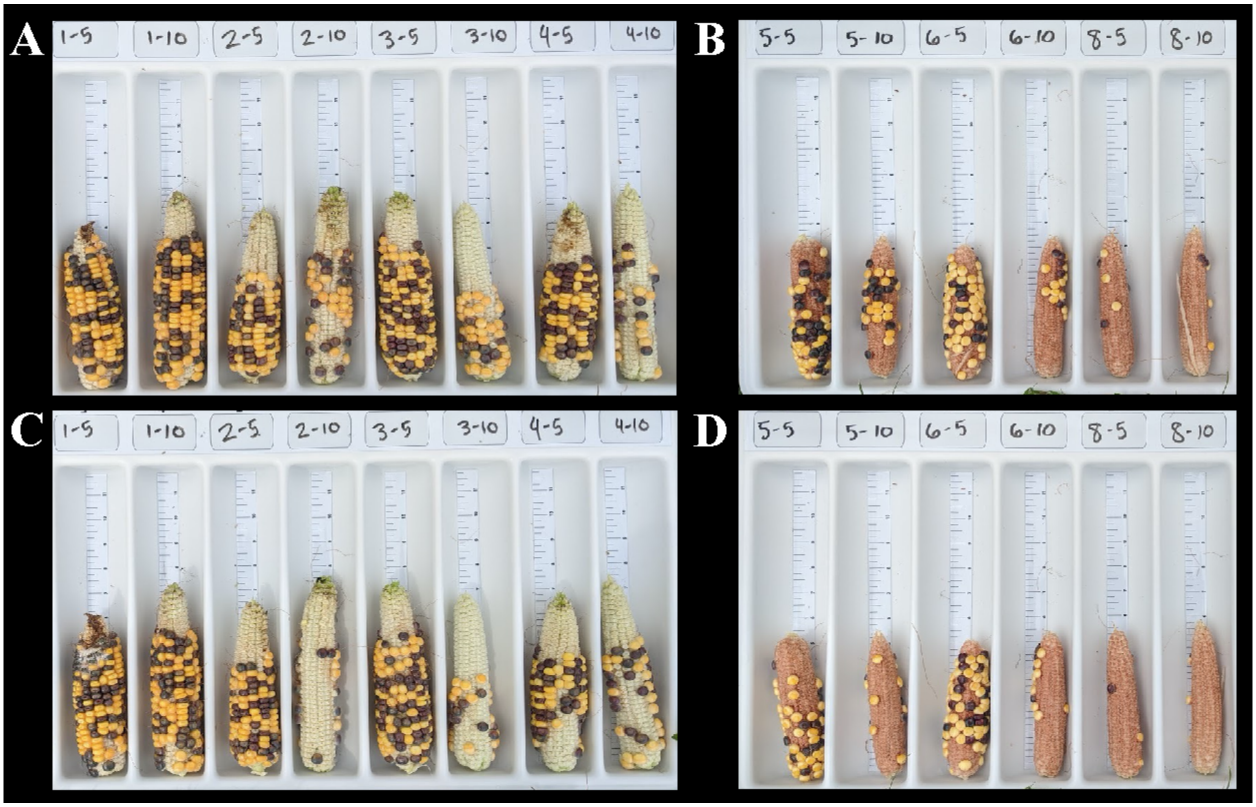

**Supplemental Figure S3:** Example maize ears at harvest pollinated with collected and stored maize pollen for A) 24 and B) 48 hours. C), D) Examples of ears at harvest pollinated using pollen collected, mixed, and directly applied to receptive silks during hybrid seed production.

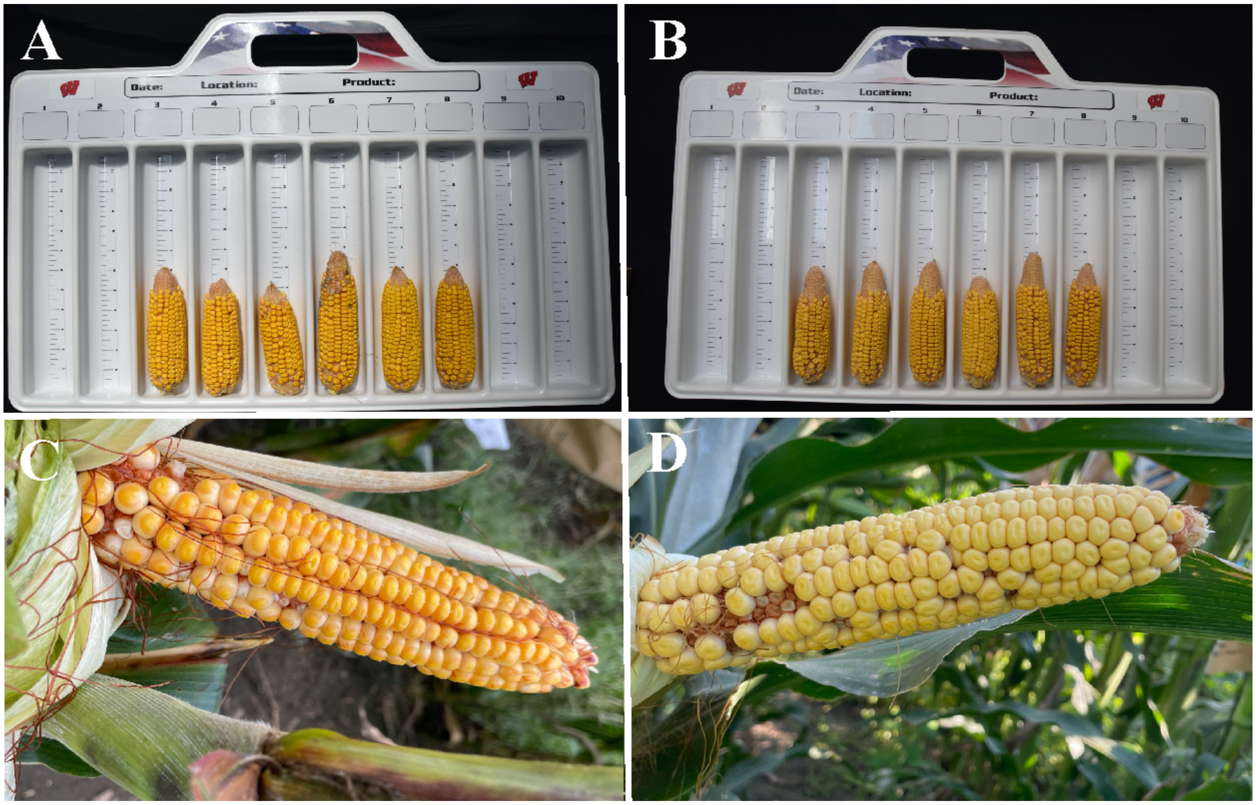

**Supplemental Table S1.** Number of kernels harvested when PHAJ0 pollen is collected and then mixed with five different substrates and stored for 2, 6, or 21 days prior to being used to pollinate two different ears (Rep 1 and Rep 2) of the inbred line LH244. The pollen was diluted to a ratio of 1:5 pollen to substrate and undiluted pollen was used as a control. '>100' shown when over 100 kernels were observed on the ears at harvest.

| | Days of Storage | | | | | |
|---|---|---|---|---|---|---|
| | **2** | | **6** | | **21** | |
| **Substrate** | Rep 1 | Rep 2 | Rep 1 | Rep 2 | Rep 1 | Rep 2 |
| Undiluted Pollen (No Substrate) | >100 | NA | 0 | 0 | 0 | 2 |
| Blue Polyethylene Microspheres (PEM) | >100 | >100 | 37 | 45 | 0 | 0 |
| Aeroperl 300/30 | 0 | 0 | 0 | 0 | 0 | 0 |
| Sipernat D 13 | 6 | 0 | 0 | 0 | 2 | 0 |
| Sipernat 22 S | 0 | 1 | 1 | 0 | 0 | 0 |
| Diatomaceous Earth | 5 | 4 | 0 | 0 | 0 | 0 |

**Supplemental Table S2.** ANOVA table describing the effect of the timing of pollen application and ratio of pollen to substrate when pollen from the inbred line PHP02 is mixed with PEEK-MP140 at a ratio of 1:5 and 1:10 pollen to substrate and stored up to 48 hours. Mixed PHP02 pollen was used to pollinate PHP02 plants with receptive silks each hour between 7:00 A.M. and 12:00 P.M.

| | **24 Hours** | | **48 Hours** | |
|---|---|---|---|---|
| | **F value** | **P-value** | **F value** | **P-value** |
| Time | 1.17 | 0.4335 | 1.35 | 0.3753 |
| Ratio | 5.37 | 0.0682 | 11.15 | 0.0206 |

**Supplemental Table S3:** Mean, maximum (Max), median, minimum (Min) and standard error (SE) in grain fill measured as the average number of kernels per ear (Count) and visually rated percent grain fill (Percent) when pollen from the inbred lines LH244, PH24E, and LH287 is collected and stored out to nine days and used to pollinate the inbred line LH244. Summary statistics shown for two different plantings dates. Blank entries represent days when no pollinations were made due to inclement weather or limited silk availability

| Planting | Inbred | Measurement | Statistic | Day 0 | Day 1 | Day 2 | Day 3 | Day 4 | Day 5 | Day 6 | Day 7 | Day 8 | Day 9 |
|---|---|---|---|---|---|---|---|---|---|---|---|---|---|
| First | LH244 | Count | Mean | 113.83 | 255.50 | 249.33 | 127.80 | 92.83 | 51.00 | 34.33 | 12.40 | 0.00 | 0.00 |
| First | LH244 | Count | Max | 198.00 | 281.00 | 332.00 | 316.00 | 150.00 | 87.00 | 75.00 | 22.00 | 0.00 | 0.00 |
| First | LH244 | Count | Median | 108.50 | 261.00 | 262.00 | 81.00 | 77.50 | 55.00 | 21.00 | 14.00 | 0.00 | 0.00 |
| First | LH244 | Count | Min | 3.00 | 209.00 | 122.00 | 35.00 | 48.00 | 8.00 | 5.00 | 3.00 | 0.00 | 0.00 |
| First | LH244 | Count | SE | 13.73 | 4.85 | 13.53 | 21.50 | 8.43 | 6.35 | 5.77 | 1.37 | | |
| First | LH244 | Percent | Max | 7.00 | 7.00 | 8.00 | 7.50 | 7.00 | 4.50 | 3.00 | 1.00 | 0.00 | 0.00 |
| First | LH244 | Percent | Median | 5.75 | 7.00 | 7.25 | 3.50 | 4.00 | 3.00 | 1.25 | 1.00 | 0.00 | 0.00 |
| First | LH244 | Percent | Min | 1.00 | 6.00 | 6.00 | 3.00 | 2.00 | 1.00 | 1.00 | 1.00 | 0.00 | 0.00 |
| First | LH244 | Percent | SE | 0.40 | 0.08 | 0.14 | 0.39 | 0.37 | 0.35 | 0.17 | 0.00 | | |
| First | LH244 | Percent | Mean | 5.08 | 6.75 | 7.25 | 4.60 | 4.25 | 2.83 | 1.67 | 1.00 | 0.00 | 0.00 |
| First | LH287 | Count | Mean | 125.00 | 20.40 | 47.14 | 64.67 | 48.20 | 11.20 | 0.00 | 3.75 | 0.00 | 3.43 |
| First | LH287 | Count | Max | 192.00 | 41.00 | 94.00 | 128.00 | 66.00 | 25.00 | 0.00 | 6.00 | 0.00 | 12.00 |
| First | LH287 | Count | Median | 127.00 | 15.00 | 50.00 | 57.50 | 46.00 | 10.00 | 0.00 | 4.00 | 0.00 | 1.00 |
| First | LH287 | Count | Min | 39.00 | 11.00 | 4.00 | 18.00 | 30.00 | 1.00 | 0.00 | 1.00 | 0.00 | 0.00 |
| First | LH287 | Count | SE | 11.97 | 2.35 | 6.11 | 6.98 | 2.79 | 1.72 | | 0.42 | | 0.79 |
| First | LH287 | Percent | Max | 7.50 | 2.00 | 5.50 | 5.50 | 3.00 | 1.50 | 0.00 | 1.00 | 0.00 | 1.00 |
| First | LH287 | Percent | Median | 7.00 | 1.50 | 2.50 | 3.00 | 2.50 | 1.00 | 0.00 | 1.00 | 0.00 | 1.00 |
| First | LH287 | Percent | Min | 3.50 | 1.00 | 1.00 | 1.00 | 2.00 | 1.00 | 0.00 | 1.00 | 0.00 | 1.00 |
| First | LH287 | Percent | SE | 0.31 | 0.08 | 0.32 | 0.30 | 0.09 | 0.04 | | 0.00 | | 0.00 |
| First | LH287 | Percent | Mean | 6.25 | 1.40 | 2.79 | 2.92 | 2.50 | 1.10 | 0.00 | 1.00 | 0.00 | 1.00 |
| First | PH24E | Count | Mean | 154.17 | 60.00 | 148.00 | 107.67 | 92.50 | 104.67 | 26.33 | 26.00 | 10.80 | 3.40 |
| First | PH24E | Count | Max | 289.00 | 98.00 | 195.00 | 166.00 | 142.00 | 178.00 | 53.00 | 50.00 | 20.00 | 9.00 |
| First | PH24E | Count | Median | 167.50 | 63.50 | 163.00 | 114.50 | 116.50 | 102.00 | 20.50 | 22.50 | 9.00 | 3.00 |

| | | | | | | | | | | | | | |
|---|---|---|---|---|---|---|---|---|---|---|---|---|---|
| First | PH24E | Count | Min | 3.00 | 29.00 | 71.00 | 24.00 | 8.00 | 30.00 | 5.00 | 14.00 | 0.00 | 0.00 |
| First | PH24E | Count | SE | 18.42 | 4.82 | 8.27 | 10.41 | 11.04 | 9.07 | 3.76 | 2.55 | 1.70 | 0.66 |
| First | PH24E | Percent | Max | 8.50 | 4.00 | 8.00 | 7.50 | 7.00 | 6.50 | 3.50 | 4.00 | 2.00 | 1.00 |
| First | PH24E | Percent | Median | 7.00 | 2.75 | 6.50 | 5.50 | 5.50 | 5.75 | 1.75 | 1.50 | 1.25 | 1.00 |
| First | PH24E | Percent | Min | 1.00 | 1.50 | 4.50 | 1.50 | 1.00 | 2.00 | 1.00 | 1.00 | 1.00 | 1.00 |
| First | PH24E | Percent | SE | 0.52 | 0.19 | 0.22 | 0.41 | 0.49 | 0.32 | 0.20 | 0.21 | 0.09 | 0.00 |
| First | PH24E | Percent | Mean | 5.92 | 2.67 | 6.42 | 4.83 | 4.50 | 5.33 | 1.92 | 1.75 | 1.38 | 1.00 |
| Second | LH244 | Count | Max | 363.00 | 298.00 | 101.00 | 64.00 | 21.00 | 14.00 | | | | |
| Second | LH244 | Count | Mean | 290.33 | 175.33 | 68.20 | 39.83 | 17.00 | 11.50 | | | | |
| Second | LH244 | Count | Median | 300.50 | 161.50 | 53.00 | 39.50 | 17.00 | 11.50 | | | | |
| Second | LH244 | Count | Min | 196.00 | 104.00 | 42.00 | 22.00 | 13.00 | 9.00 | | | | |
| Second | LH244 | Count | SE | 10.37 | 13.04 | 5.53 | 3.08 | 1.07 | 0.67 | | | | |
| Second | LH244 | Percent | Max | 8.00 | 8.50 | 4.00 | 2.50 | 1.00 | 1.00 | | | | |
| Second | LH244 | Percent | Mean | 7.67 | 6.08 | 2.70 | 1.67 | 1.00 | 1.00 | | | | |
| Second | LH244 | Percent | Median | 8.00 | 6.50 | 2.50 | 1.75 | 1.00 | 1.00 | | | | |
| Second | LH244 | Percent | Min | 7.00 | 3.00 | 1.50 | 1.00 | 1.00 | 1.00 | | | | |
| Second | LH244 | Percent | SE | 0.10 | 0.38 | 0.20 | 0.11 | 0.00 | 0.00 | | | | |
| Second | LH287 | Count | Max | 265.00 | 252.00 | 99.00 | 133.00 | 27.00 | 1.00 | 2.00 | 0.00 | | |
| Second | LH287 | Count | Mean | 142.17 | 103.83 | 66.83 | 50.67 | 9.83 | 0.50 | 1.33 | 0.00 | | |
| Second | LH287 | Count | Median | 127.50 | 80.00 | 68.50 | 44.00 | 6.50 | 0.50 | 1.00 | 0.00 | | |
| Second | LH287 | Count | Min | 40.00 | 30.00 | 31.00 | 5.00 | 2.00 | 0.00 | 1.00 | 0.00 | | |
| Second | LH287 | Count | SE | 16.97 | 15.55 | 5.27 | 8.32 | 1.78 | 0.13 | 0.11 | | | |
| Second | LH287 | Percent | Max | 8.00 | 8.00 | 5.00 | 5.50 | 2.50 | 0.50 | 0.50 | 0.50 | | |
| Second | LH287 | Percent | Mean | 5.75 | 4.33 | 3.33 | 2.58 | 1.17 | 0.50 | 0.50 | 0.50 | | |
| Second | LH287 | Percent | Median | 6.00 | 4.25 | 3.50 | 2.25 | 1.00 | 0.50 | 0.50 | 0.50 | | |
| Second | LH287 | Percent | Min | 3.00 | 1.00 | 1.50 | 1.00 | 0.50 | 0.50 | 0.50 | 0.50 | | |
| Second | LH287 | Percent | SE | 0.37 | 0.51 | 0.32 | 0.30 | 0.13 | 0.00 | 0.00 | | | |
| Second | PH24E | Count | Max | 379.00 | 406.00 | 404.00 | 276.00 | 232.00 | 80.00 | 24.00 | | | |

| | | | | | | | | | | |
|---|---|---|---|---|---|---|---|---|---|---|
| Second | PH24E | Count | Mean | 335.83 | 295.67 | 272.00 | 160.83 | 89.17 | 47.83 | 17.75 |
| Second | PH24E | Count | Median | 345.00 | 304.00 | 283.50 | 147.00 | 71.50 | 50.50 | 19.00 |
| Second | PH24E | Count | Min | 293.00 | 198.00 | 96.00 | 85.00 | 8.00 | 14.00 | 9.00 |
| Second | PH24E | Count | SE | 6.70 | 14.87 | 21.18 | 12.94 | 16.47 | 5.27 | 1.34 |
| Second | PH24E | Percent | Max | 8.50 | 9.00 | 9.00 | 7.00 | 7.00 | 4.50 | 1.50 |
| Second | PH24E | Percent | Mean | 8.00 | 8.00 | 7.50 | 5.42 | 3.50 | 2.50 | 1.25 |
| Second | PH24E | Percent | Median | 8.00 | 8.25 | 7.50 | 5.50 | 3.25 | 2.75 | 1.25 |
| Second | PH24E | Percent | Min | 7.50 | 7.00 | 6.00 | 3.50 | 1.00 | 1.00 | 1.00 |
| Second | PH24E | Percent | SE | 0.06 | 0.16 | 0.19 | 0.21 | 0.50 | 0.25 | 0.05 |
